## Supplementary Information for "A deeply conserved protease, acylamino acid-releasing enzyme (AARE), acts in ageing in Physcomitrella and Arabidopsis"

### Supplementary Figures

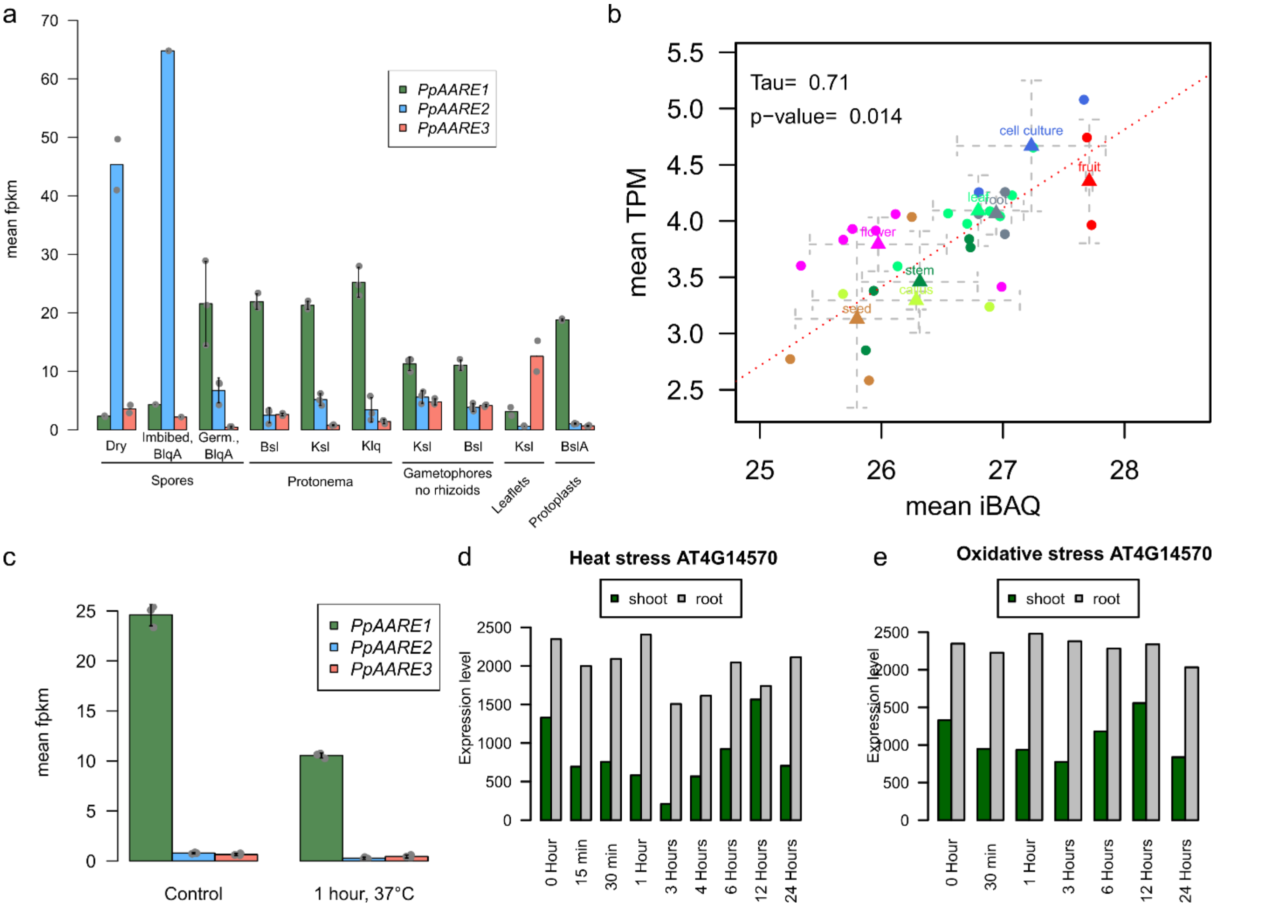


**Fig. S1 Expression levels of *PpAARE* genes and *AtAARE* in different tissues and under stress conditions.** (a) Expression levels of the three *PpAARE* genes in different tissue and cell culture types. All data have been downloaded from *PEATmoss* (https://peatmoss.online.uni-marburg.de/) and are described in [1,2]. Depicted are mean FPKM (Fragments Per Kilobase Million) values with standard deviation. Data points represent biological replicates, in some cases only data from two replicates (dry spores, leaflets (Ksl)) or only from one (Imbibed spores) replicate were available. . Abbreviations: B = BCD medium[3]; lq = liquid; A =ammonium tartrate; sl = solid; K = Knop medium; Germ. = germinating. (b) Tissue-specific expression levels of *AtAARE* and correlation between protein abundance and transcript level in Columbia WT (Col0). Transcript and protein abundance data[4] have been downloaded from the *ATHENA* database (http://athena.proteomics.wzw.tum.de/). Transcript levels are represented by transcripts per million kilobase (TPM) and protein abundances are represented as intensity based absolute quantification values (iBAQ). Triangles are mean values with standard deviation (dashed lines) for each tissue group. Individual data points of each category are plotted as filled circles.

Pearson´s correlation ($r = 0.85$) indicates a strong positive and significant ($p\text{-value} = 0.007$) correlation of expression levels and protein abundance in major tissue types. The correlation was calculated based on the means of the tissue groups. (c) Expression levels for *PpAARE1*, *PpAARE2* and *PpAARE3* in protonema in response to heat stress. Depicted are mean FPKM values from three biological replicates with standard deviation. Data were downloaded from PEATmoss (https://peatmoss.online.uni-marburg.de/) and are described in [1,2]. (d) Expression levels of *AtAARE* in response to heat and oxidative stress (e). Heat stress (38°C) was applied for 3 hours[5], samples were taken after recovery at 25°C at indicated timepoints. Bars represent the mean of two biological replicates deviation. All data (d, e) presented were downloaded from the Arabidopsis eFP Browser (http://bar.utoronto.ca/efp_arabidopsis/)[5,6].

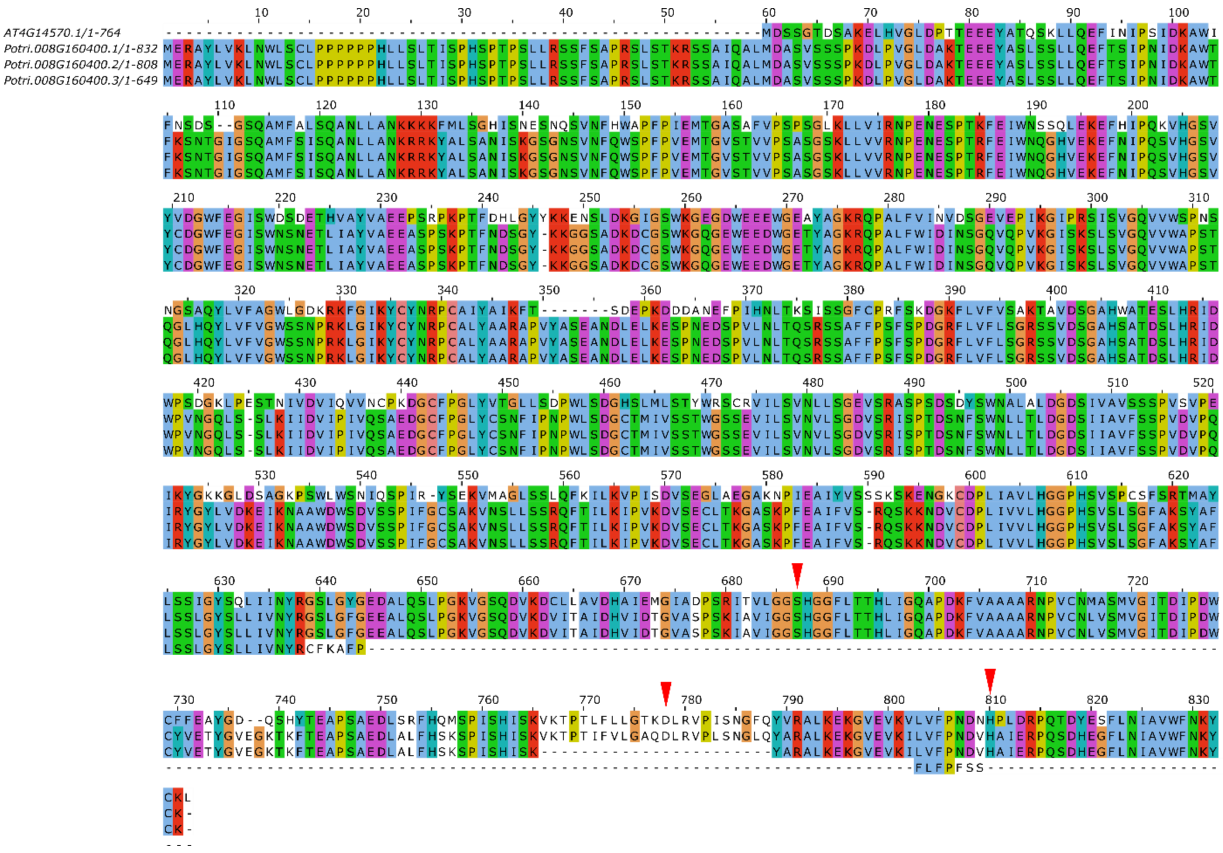

**Fig. S2 Sequence alignment of AtAARE and PtAARE isoforms.** Red arrows point to the catalytic residues (Ser, Asp, His) of AtAARE as described[7]. Two splice variants result in non-functional protein isoforms. Potri.008G160400.2 lacks the catalytic Asp (alignment position 778) and Potri.008G160400.3 is lacking the whole catalytic domain. Sequences were aligned in Jalview[8] (Version: 2.11.1.3) using T-Coffee[9] with default settings.

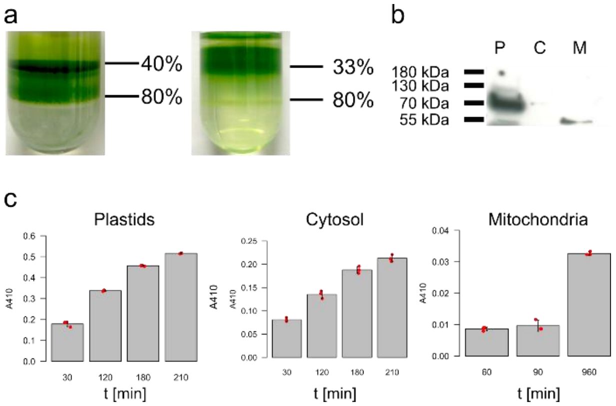


**Fig. S3 AARE activity in Physcomitrella plastids and mitochondria.** (a) Plastids and mitochondria were simultaneously isolated from the protonema as described[10]. Intact plastids were collected at the interface between 80% and 40% Percoll, mitochondria were collected at the interface between 80% and 33% Percoll. (b) Cross-contamination of the obtained fractions with plastid proteins was assessed via Western Blot against the plastid HSP70B (Agrisera, AS06 175). Fractions were soluble plastid proteins (P), the soluble fraction depleted of plastids and mitochondria (referred to as cytosol, (C)) and soluble mitochondrial proteins (M). The intracellular fraction was free of plastid cross-contaminations. A full blot image is available in Fig. S10. (c) AARE activity measured using AcAla-pNA is present in isolated plastid fractions and in organelle-depleted intracellular fractions. Only low activity is detectable in mitochondrial fractions. The activity is indicated by the increase of the absorbance at 410 nm ($A_{410}$) over time caused by the release of pNA. Mean values from technical triplicates with standard deviation are shown.

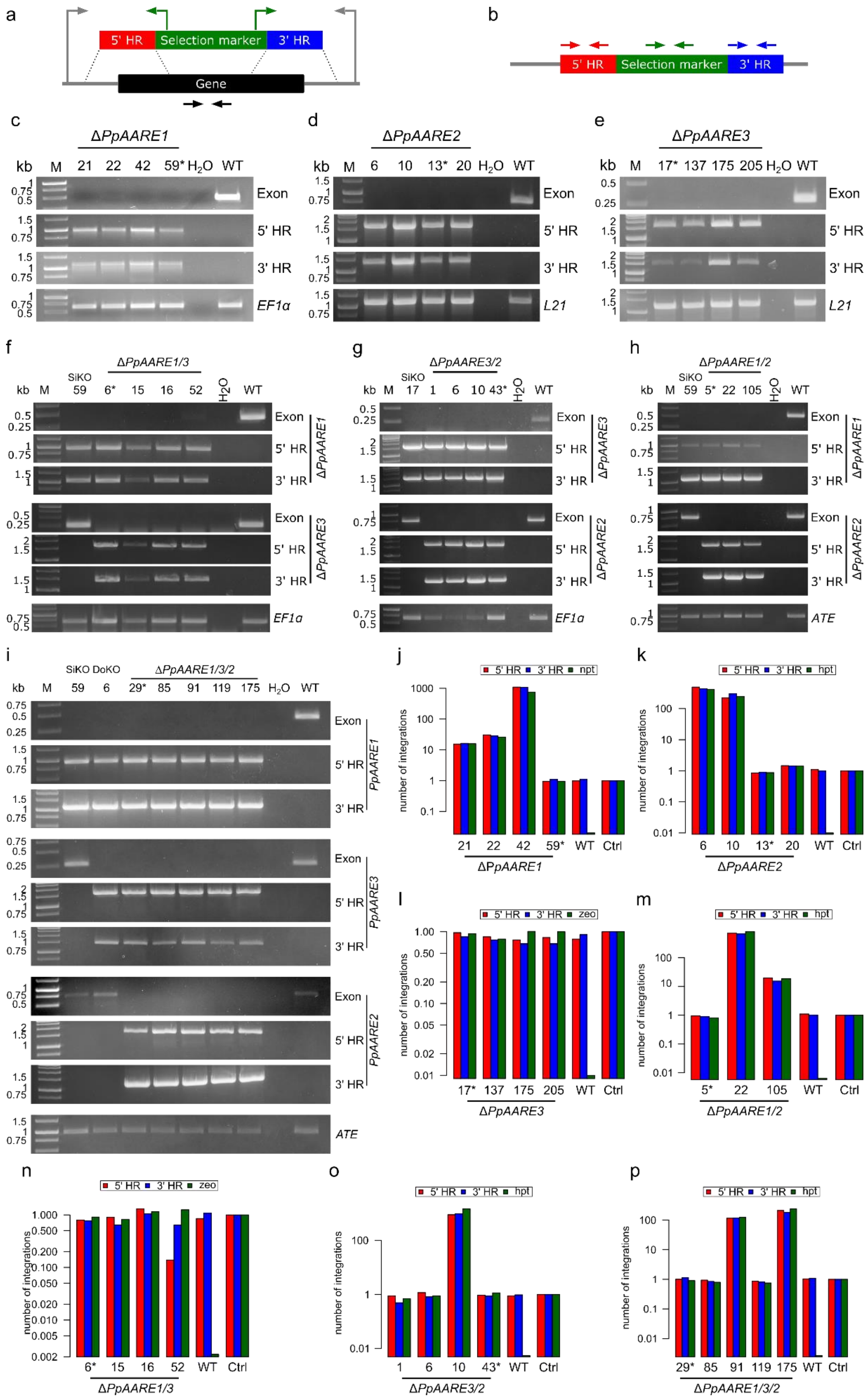

a
5' HR
Selection marker
3' HR
Gene
b
5' HR
Selection marker
3' HR
c
ΔPpAARE1
kb M 21 22 42 59* H2O WT
Exon
5' HR
3' HR
EF1α
d
ΔPpAARE2
kb M 6 10 13* 20 H2O WT
Exon
5' HR
3' HR
L21
e
ΔPpAARE3
kb M 17* 137 175 205 H2O WT
Exon
5' HR
3' HR
L21
f
ΔPpAARE1/3
SiKO
kb M 59 6* 15 16 52 H2O WT
Exon
5' HR
3' HR
ΔPpAARE1
ΔPpAARE3
EF1α
g
ΔPpAARE3/2
SiKO
kb M 17 1 6 10 43* H2O WT
ΔPpAARE3
ΔPpAARE2
EF1α
h
ΔPpAARE1/2
SiKO
kb M 59 5* 22 105 H2O WT
ΔPpAARE1
ΔPpAARE2
ATE
i
SiKO DoKO
ΔPpAARE1/3/2
kb M 59 6 29* 85 91 119 175 H2O WT
PpAARE1
PpAARE3
PpAARE2
ATE
j
5' HR 3' HR npt
number of integrations
21 22 42 59* WT Ctrl
ΔPpAARE1
k
5' HR 3' HR hpt
6 10 13* 20 WT Ctrl
ΔPpAARE2
l
5' HR 3' HR zeo
17* 137 175 205 WT Ctrl
ΔPpAARE3
m
5' HR 3' HR hpt
5* 22 105 WT Ctrl
ΔPpAARE1/2
n
5' HR 3' HR zeo
6* 15 16 52 WT Ctrl
ΔPpAARE1/3
o
5' HR 3' HR hpt
1 6 10 43* WT Ctrl
ΔPpAARE3/2
p
5' HR 3' HR hpt
29* 85 91 119 175 WT Ctrl
ΔPpAARE1/3/2

**Fig. S4 Identification of PpAARE single, double and triple KO mutants by PCR on genomic DNA.** (a) PCR scheme showing the primer combinations (arrows) used to validate KOs. One pair each is used to check for the correct positioning of the homologous flanks (5' and 3' HR) at the desired locus (green, grey) and one pair was used to check for the presence/absence of the genomic region (black arrows). (b) Scheme showing primers (arrows) used for qPCR on genomic DNA to determine the copy numbers of the KO construct integrated in the genome. Primers are listed in Supplementary Table S2 and Supplementary Table S5. Identification of single KOs for *PpAARE1* (c), *PpAARE2* (d) and *PpAARE3* (e). Identification of Δ*PpAARE1/3* (f), Δ*PpAARE3/2* (g), Δ*PpAARE1/2* (h) double KO lines and Δ*PpAARE1/3/2* triple KO lines (i). SiKO indicate the single KO line employed as parental line for the generation of a double KO. Δ*PpAARE1/3* #6 was employed as parental line for the generation of Δ*PpAARE1/3/2* triple KO lines. Expected amplicon sizes for *PpAARE1* KO PCR: Exon: 497 bp; 5' HR: 983 bp; 3' HR: 1128 bp. Expected amplicon sizes for *PpAARE2* KO PCR: Exon: 768 bp; 5' HR: 1678 bp; 3' HR: 1309 bp. Expected amplicon sizes for *PpAARE3* KO PCR: Exon: 273 bp; 5' HR: 1631 bp; 3' HR: 1486 bp. *EF1α*, *L21* and ATE are reference genes (Supplementary Table S2). Expected amplicon sizes: *EF1α*: 660 bp; *L21*: 1173 bp; 970 bp. (j)-(p) Copy number count of the respective KO constructs for each validated KO line determined via qPCR. Ctrl: Single copy integration control for each employed selection marker. Stars indicate lines with only a single genomic integration of the KO construct selected for further experiments.

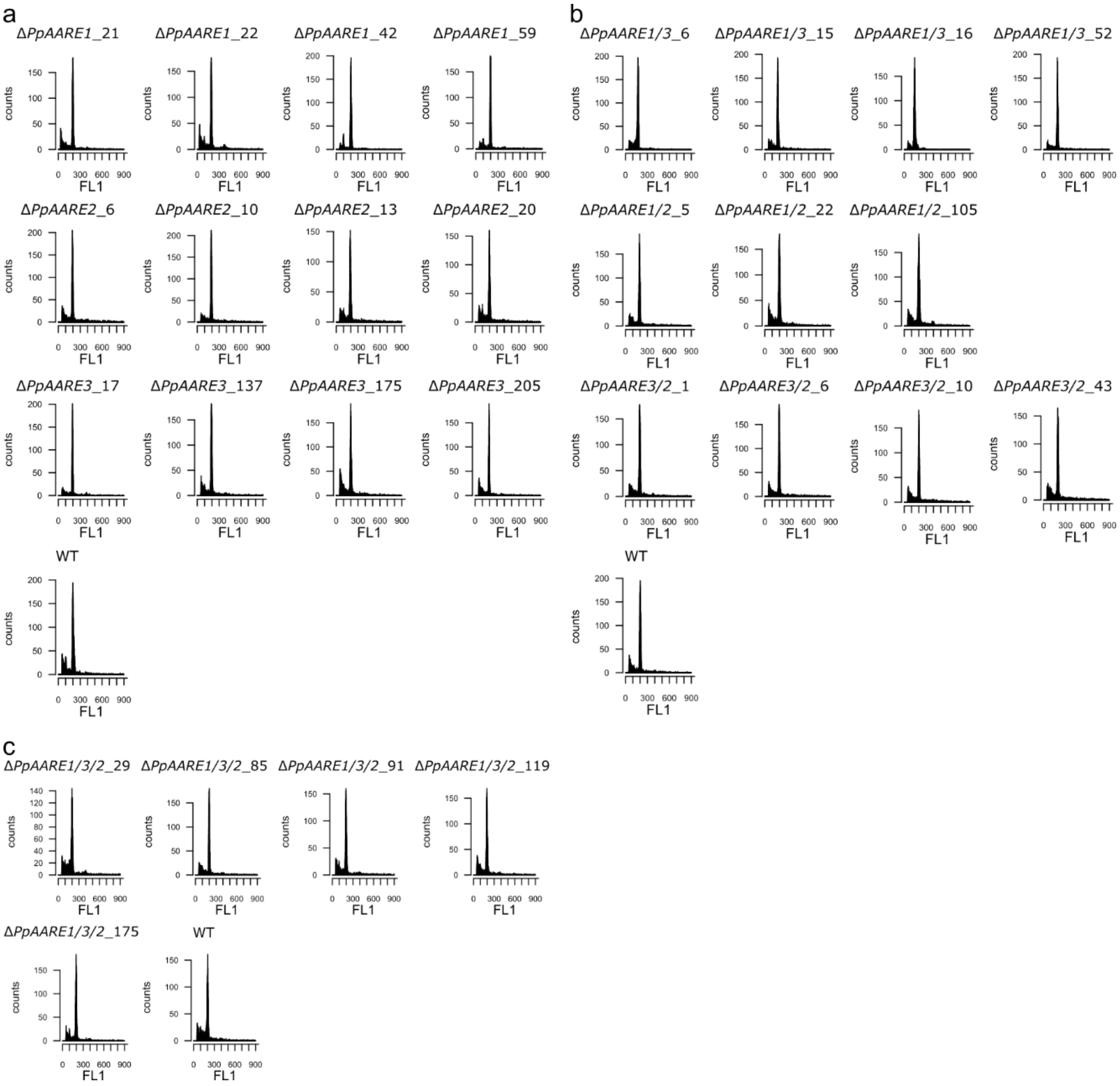


**Fig. S5 FCM analysis of all confirmed PpAARE KO lines to analyse the ploidy.** (a) Single KO lines, (b) double KO Lines, (c) triple KO lines. All lines have the major signal at ~200 like the haploid wild type. Ploidy was determined in protonema as described[11].

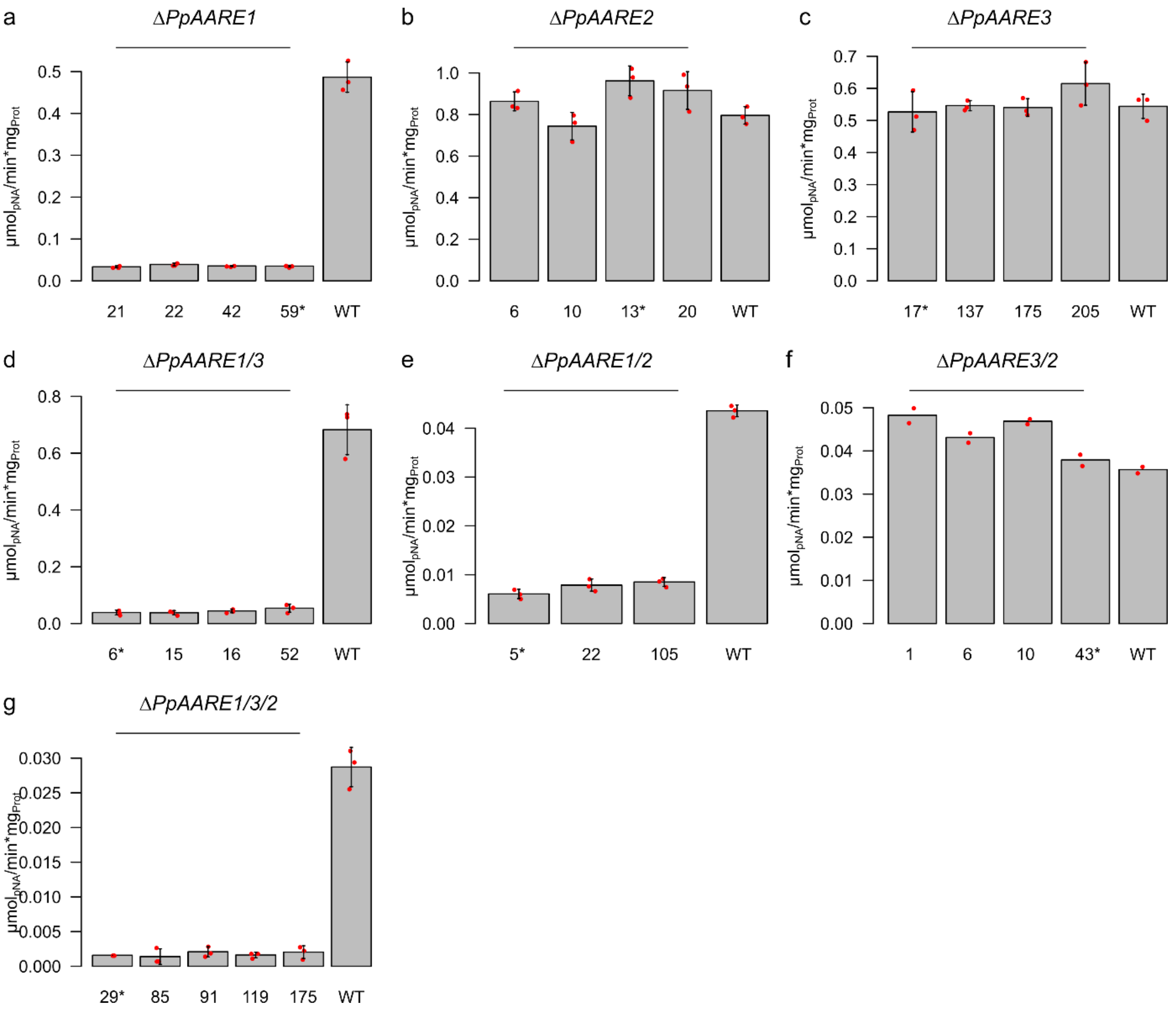


**Fig. S6 AARE exopeptidase activity in protonema of KO mutants.** AcAla-pNA was used as substrate and activity was measured in protonema. Depicted is the mean activity of three technical replicates with standard deviation. Means of activity in Δ*PpAARE3/2* lines are based on technical duplicates. All lines with a knockout of *PpAARE1* have a strongly reduced activity (a, d, e, and g). The mean activity is not reduced in Δ*PpAARE2* (b) and Δ*PpAARE3* lines (c). The double knockout of Δ*PpAARE3/2* (f) results in a slight increase of activity in lines #1, #6, #10. Stars at the line numbers indicate the lines with only a single copy of the respective knockout construct which were used for the major experiments in this study.

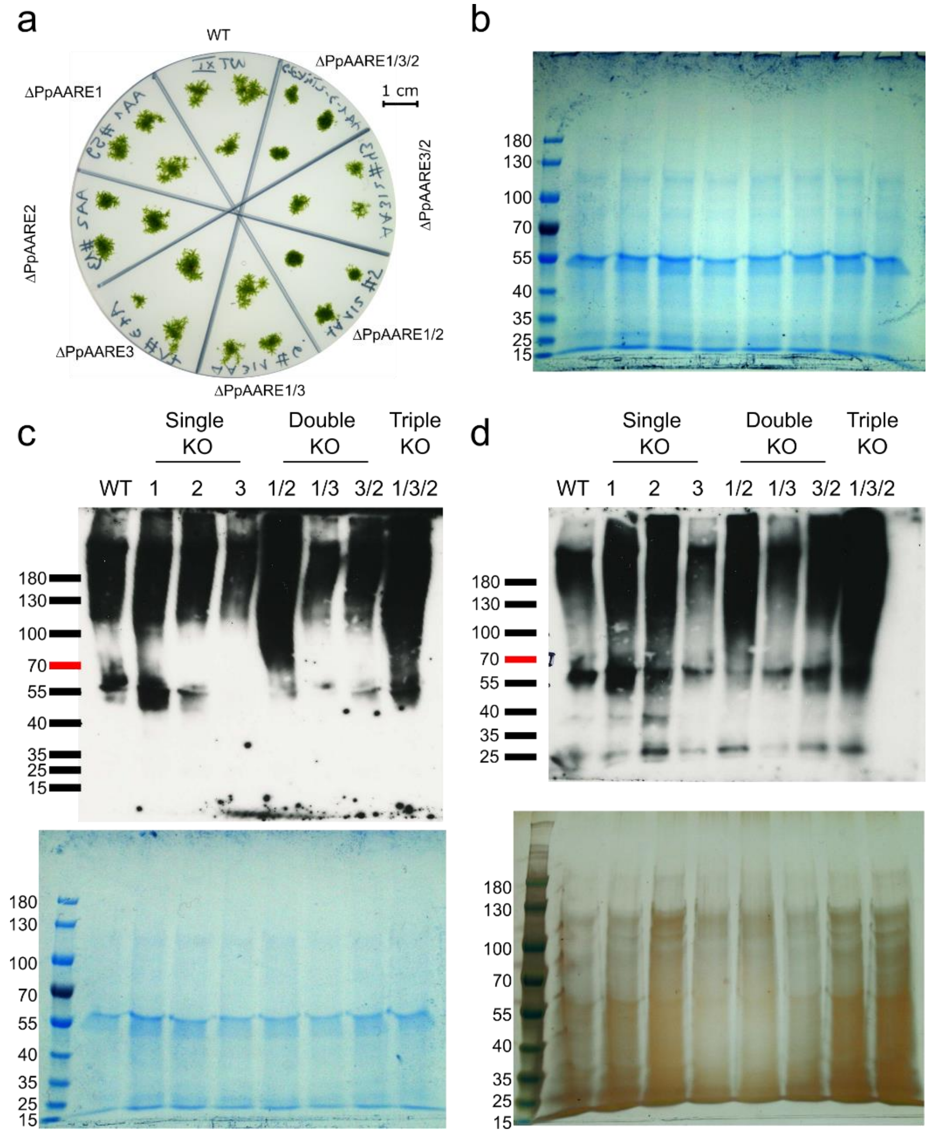


**Fig. S7 Plate overview and additional OxyBlots from Physcomitrella gametophore colonies with loading controls.** (a) Exemplary plate (KnopME) with five-week-old gametophore colonies from WT, single, double and triple knockouts. Protein extracts from single colonies were used for OxyBlot analysis. (b) Coomassie-stained loading control for the OxyBlot depicted in Fig. 3. (c+d) Additional OxyBlots with Coomassie- (c) and silver-stained (d) loading controls. Equal volumes from the same derivatized protein solutions were loaded for the OxyBlot as well as for the loading control gels. PageRuler™ Pre-stained Protein Ladder (Thermo Scientific™, #26166) was used for all Blots and loading control gels. OxyBlot analysis was performed with the OxyBlot™ *Protein Oxidation Detection Kit* (Merck) according to the manufacturer's instructions.

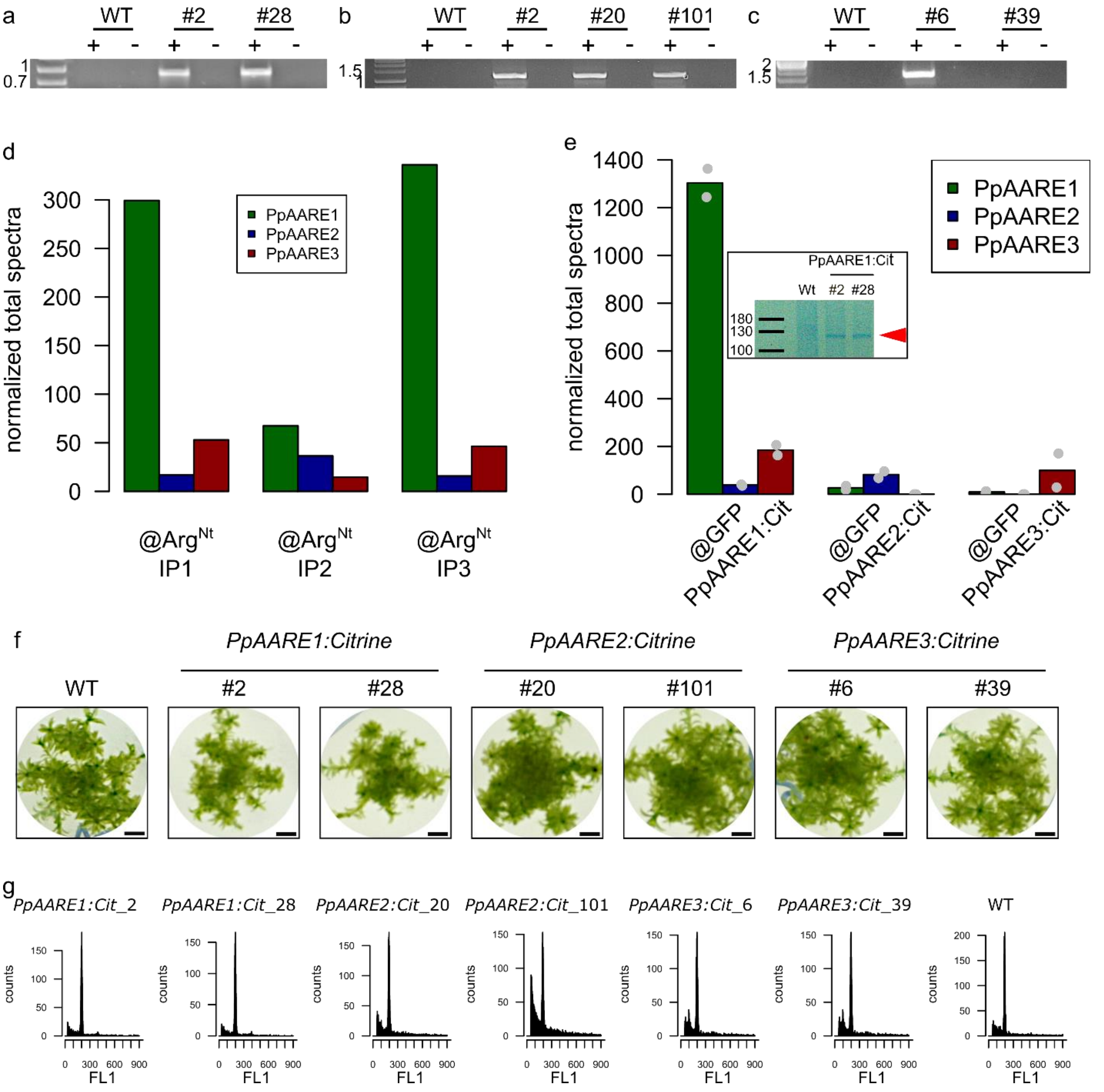


**Fig. S8 Screening for haploid Physcomitrella lines with a citrine fusion transcript and overview on identification of the three PpAARE isoforms in different IP experiments.** (a)-(c): RT PCR on citrine fusion candidate lines for *PpAARE1* (a), *PpAARE2* (b), and *PpAARE3* (c). +: with cDNA template; -: without reverse transcriptase. For each PCR a forward primer binding in the CDS of the respective PpAARE isoform and a reverse primer binding in the CDS of citrine was chosen. Primers are listed in Supplementary Table S2. No transcript was detected in *PpAARE3:Citrine* #6. *PpAARE2:Citrine* #2 was not used for further experiments. (d) All PpAARE isoforms were identified in the IP experiments targeting N-terminal arginylation (PXD003232[12,13]).

Identified proteins are listed in Supplementary Table S7. The bars represent single replicate values. (e) Overview on identification of PpAARE isoforms in the test IP experiments targeting the citrine tag of the different fusion proteins. The test IP against PpAARE1:Citrine was performed using *Dynabeeds M270 Epoxy Co-Immunoprecipitation Kit* (Life Technologies, Carlsbad, USA) with a monoclonal anti-GFP antibody (Roche, REF 11 814 460 001) as described[12] for two independent fusion lines (#2 and #28). Eluted proteins were separated via SDS-PAGE and bands (inset, red arrow) corresponding to the expected size (112 kDa) were excised and analysed by mass spectrometry. Test IP experiments against citrine-tagged PpAARE2 and PpAARE3 were performed with GFP-Trap Magnetic Particles (Chromotek) followed by an on-bead digestion with trypsin and subsequent mass spectrometry. Values represent means normalized total spectra obtained in each two independent fusion lines (PpAARE1: #2, #28; PpAARE2: #20, #101; PpAARE3: #6, #39) The mass spectrometry proteomics data have been deposited at the ProteomeXchange Consortium via the PRIDE partner repository[14,15] with the dataset identifier 10.6019/PXD038742. (f) Gametophore colonies of selected Citrine fusion lines cultivated 4 months on solid medium. Bar = 2 mm. (g) FCM analysis of all confirmed PpAARE citrine fusion lines. Ploidy was determined in protonema as described[11]. All lines have the major signal at ~200 like the haploid wild type.

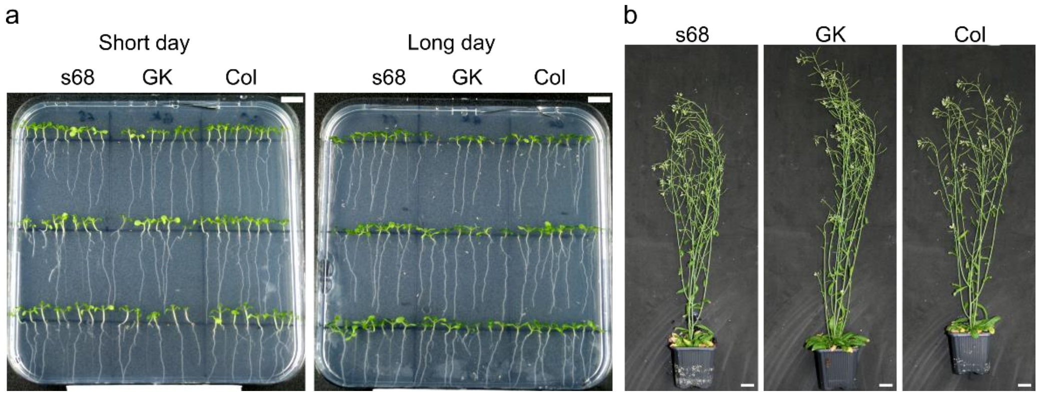


**Fig. S9 Phenotypes of seedlings and adult AtAARE T-DNA mutant lines (GK, s68) and WT (Col).** (a) AtAARE T-DNA mutant lines and WT germinated in short- or long-day conditions. Short day: 8h light /16h dark, 35 µmol/m$^2$s; Long day: 16h light / 8h dark, 35 µmol/m$^{-2}$s. Bars correspond to 1 cm (b) Adult plants cultivated for 7 weeks. Plants were grown under short day conditions (8h light/16h dark) for 4 weeks and then transferred to long day conditions for another 3 weeks (16h light/8h dark). Bars correspond to 2 cm.

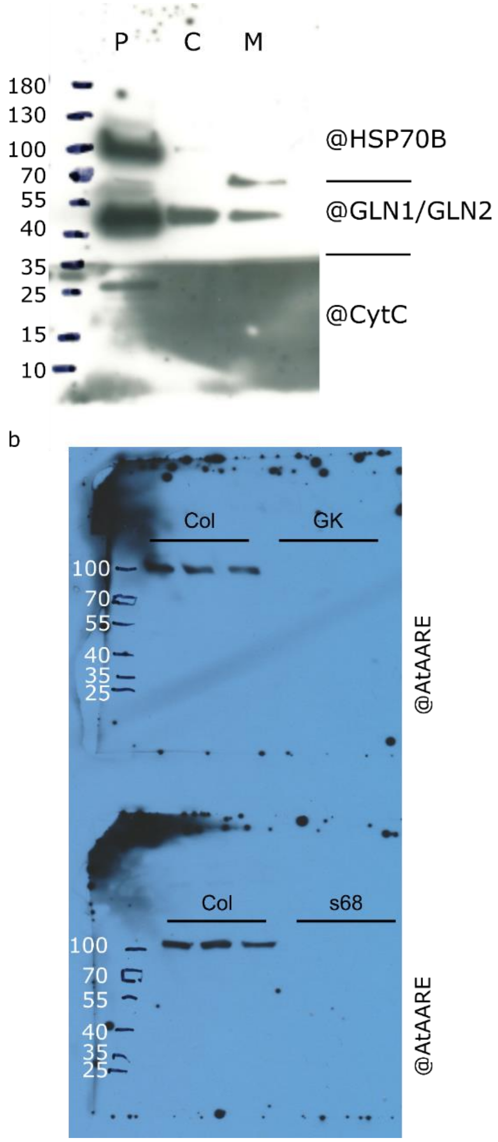


**Fig. S10 Full Western blot images of blots depicted in Fig S3b and Fig 6b.** Uncropped image of the Western blot depicted in Fig. S3b (a). The blotted membrane was cut at the indicated lines and membrane parts were incubated with three different antibodies: @HSP70B (anti heat shock protein 70B, Agrisera, AS 06 175), @GLN1/GLN2 (anti-Glutamine synthase, Agrisera, AS 08 295) and @CytC (anti-Cytochrome C, Agrisera, AS 08 343). Expected signal sizes were ~72 kDa for @HSP70B, ~40 kDa for @GLN1/GLN2 and ~25 kDa for @CytC. Uncropped image of the @AtAARE Western blot depicted in Fig 6b (b). The antibody was kindly provided by Dr. Yasuo Yamauchi and is described in [16].
